# Aligning conservation status, vulnerability factors, and ecological and evolutionary uniqueness to produce integrated assessments of the world’s birds

**DOI:** 10.1101/2025.07.15.664934

**Authors:** Eliot T. Miller, Jeffery L. Larkin, Anna M. Matthews, Michael Parr, James J. Giocomo, Daniel J. Lebbin

**Affiliations:** American Bird Conservancy, The Plains, VA 20198, USA; Department of Biology, Indiana University of Pennsylvania, Indiana, PA 15705, USA

**Keywords:** Biodiversity monitoring, Birds, Ecological integrity metrics, Conservation prioritization, Phylogenetic and functional uniqueness, Global biodiversity assessments

## Abstract

A growing awareness, now enshrined in the Kunming-Montreal Global Biodiversity Framework, of the need to monitor biodiversity effectively at scale has led to a proliferation of novel solutions for doing so. Although global biodiversity encompasses all life, from tiny nitrogen-fixing bacteria to emergent rainforest trees, birds have several characteristics that make them a frequent focus of such monitoring efforts. In particular, birds frequently give diagnostic, species-specific vocalizations that simplify monitoring, they perform a number of critical ecosystem services, they are widely distributed in most ecosystems with strong representation on all continents, and the basic ecology, conservation status, populations, and distributions of many species is well known; birds thus provide a window into the underlying health and habitats of the systems under study. How best to summarize biodiversity monitoring results is a research question that has led to the development of approaches that incorporate species’ IUCN Red List threat assessments into site-level biodiversity scores. Notably, birds’ vocal behavior means that they can be effectively surveyed at scale with passive acoustic monitoring, and the potential to link such monitoring with automated identification and therefore quickly generate site-level biodiversity scores is an appealing approach to implement rigorous evaluations of global biodiversity. Yet, while many of the world’s birds are suffering worrisome population declines, the vast majority of species (78%) are still ranked “Least Concern” by the Red List. In an effort to develop a species scoring system that would be more conducive to such site-level valuations, we integrated key databases of species’ population status assessments, exposure to known vulnerability factors, and their functional and phylogenetic uniqueness to provide quantitative summaries of their conservation significance. We augmented these databases with two novel data sets available for most of the world’s birds: quantitative measurements of migration distances, and species-level phylogenetic and functional uniqueness values comparing each species to those it co-occurs with throughout its range. While the resulting BirdsPlus species scores also inherently reflect our own scientific expertise and judgement, our approach is transparent, dynamic, easily updated, and readily modified by users with different goals or values.

## INTRODUCTION

The IUCN (International Union for the Conservation of Nature) Red List (IUCN 2022) has, for decades, helped to identify species and geographic areas in need of urgent conservation action. First released in 1966 (Simon 1966), the frequency with which the Red List is used in conservation decisions, the regular updates, and the global attention focused on the list attest to the critical need for such population assessments (Butchart et al. 2025). The Red List can help prioritize conservation of the most endangered species, or on sites that host such species. Setting aside and managing protected areas in this way is essential for biodiversity conservation (Luther et al. 2021), and underpins major global conservation initiatives such as Target 3 of the Kunming-Montreal Global Biodiversity Framework (the “30×30 target” (Convention on Biological Diversity 2022)). However, conserving nature outside protected areas through the implementation of conservation practices on working lands is also critical (Ciuzio et al. 2013; Bennett et al. 2018; Michel et al. 2020; Dertien & Baldwin 2022).

One good example of scalable working lands conservation efforts that can be utilized to benefit wildlife and biodiversity are programs and initiatives offered by the United States Department of Agriculture-Natural Resources Conservation Service. For example, the agency’s Working Lands for Wildlife and Regional Conservation Partnership Programs focus on restoring and enhancing habitat for target species such as the Golden-winged Warbler, *Vermivora chrysoptera* (McNeil et al. 2020), Cerulean Warbler, *Setophaga cerulea* (Shaffer et al. 2025), Greater Sage Grouse, *Centrocercus urophasianus* (Naugle et al. 2024), and Northern Bobwhite, *Colinus virginianus* (Burger Jr. et al. 2006). Several successful state-led and partnership-led wildlife conservation programs that mimic aspects of the NRCS-Environmental Quality Incentive Program have also been developed in the past 10-15 years, such as the Oaks and Prairies Joint Venture’s Grassland Restoration Incentive Program (GRIP) which has been adopted by several Migratory Bird Joint Ventures (Giocomo et al. 2017). Additionally, many conservation groups, including American Bird Conservancy (“Bird-Friendly Working Lands” n.d.), National Audubon Society (“Conservation Ranching | Audubon” 2025), BirdLife International (“Biodiversity -Grassland Alliance” n.d.), and the Smithsonian Institution (“Virginia Working Landscapes” 2025) have promoted conservation practices on farms and ranches to enhance habitat for birds. One obstacle to scaling this work is the need for metrics that can quantify the conservation benefits after management intervention. Monitoring is critical to gauging program effectiveness and modifying practice implementation. Easy-to-interpret metrics that move beyond simple estimates of species occurrence are particularly desirable, since they could catalyze further conservation investment, including from non-philanthropic sources (Rodewald et al. 2020). While there is a rich suite of literature and countless cases employing traditional sources of monitoring such as distance sampling (Buckland et al. 2001), many researchers, companies and NGOs are working to develop tools to provide biodiversity metrics that cost-effectively link monitoring at scale with emerging technologies such as LiDAR (McNeil et al. 2023), eDNA (Beng & Corlett 2020), camera traps (Cordier et al. 2022), and passive acoustic monitoring paired with machine learning classifiers like BirdNET (Wood et al. 2022; Larkin et al. 2024). There is great potential in these technologies, and while such approaches often incorporate an element of the conservation significance of detected species, the default has been to use coarse threat assessments to guide these valuations (Mair et al. 2021).

From a quantitative and practical perspective, however, coarse, ordinal assessments like the Red List neither offer the resolution, nor express the diversity of relevant conservation considerations that some practitioners desire. Often, more nuanced assessments are available, but their regionally limited nature can hamper efforts to extend these approaches to larger spatial scales (Michel et al. 2020). Species-level factors that have emerged as valuable indicators of conservation priority, beyond threat status, include: ecological function (Díaz et al. 2013; Asner et al. 2017), phylogenetic uniqueness (Faith 1992), and the identification of characteristics that can predispose species to adverse outcomes, such as long-distance migration (Saunders et al. 2025) or human exploitation (Ripple et al. 2016). While some indices have endeavored to incorporate elements of these factors, e.g., phylogenetic uniqueness (Isaac et al. 2007), none have sought to derive a comprehensive, integrated conservation score that incorporates a wide variety of such factors. Moreover, while the IUCN Red List is globally authoritative, other, regionally or taxonomically limited population assessments are available that provide a more fine-grained understanding of population status and trends (Fink et al. 2021; SoIB 2023; Kittelberger et al. 2023; Partners in Flight 2024). Harmonizing these disparate data sources has the potential to produce a comprehensive conservation score at the global level that incorporates not only other key factors of potential relevance, but more detailed population trends when available.

Critical features to include in generating a conservation valuation score are that it should be data-driven, easy to update, and values must be available for all species in question (Burgess et al. 2024). Moreover, it should incorporate species’ characteristics that the research community has identified as important, such as phylogenetic and functional uniqueness (Eisenhauer et al. 2023). While no score will ever be perfect, and there is inherent subjectivity in such an exercise, data-driven approaches can be more objective, easier to modify and update, and greatly facilitate comparison between sites. Birds, in particular, are well-suited to such an approach, owing to their rapid responses to environmental change and habitat specificity (Stillman et al. 2025), well-resolved taxonomy (Rheindt et al. 2025) and phylogeny (McTavish et al. 2025), and the long-standing, substantial contributions of citizen scientists to understanding their distributions at a global scale (Sullivan et al. 2014). We used these convenient properties to develop a conservation score for the world’s birds that is intended to value more than just species richness. The factors used to generate these scores include three broad categories that are combined to produce a final value: conservation status, vulnerability to known threats, and ecological and phylogenetic uniqueness. We call these values BirdsPlus species scores. The name is meant to indicate that biodiversity is multifaceted; we focus on birds as indicators, and we include elements beyond species richness, with the intention to help quantify ecological integrity and restoration progress, amongst other things. Here we outline the methods taken to create these scores, and we make the scores and associated computer code freely available to the conservation community.

## METHODS

### Source of phylogeny

A key resource we used to create BirdsPlus scores is a comprehensive global avian phylogeny. The phylogeny and its underlying taxonomy served three critical functions: facilitating the alignment and integration of disparate datasets, calculation of phylogenetic uniqueness indices, and phylogenetic imputation of missing data. We used the *clootl* package (https://github.com/eliotmiller/clootl) to access the most recently released global avian phylogeny (McTavish et al. 2025). This is v1.4 of the phylogeny, and we output it in the 2023 Clements taxonomy, which includes 11,017 species-level bird taxa.

### Source of conservation assessments

We incorporated three sources of conservation assessment data. First, we used the IUCN Red List, which uses BirdLife taxonomy (Burfield et al. 2017). Thus, to assemble these scores, we first used IUCN scores that had previously been mapped to the 2023 Clements taxonomy (Birds of the World, downloaded on 31 May 2024). Due to taxonomic mismatches, 810 species in Clements are considered as unassessed by the IUCN. These mismatches tend to be fairly subtle issues related to taxon concepts (Lepage et al. 2014). For example, American Robin (*Turdus migratorius*, including subspecies *confinis*) does not have an IUCN status in Clements taxonomy because BirdLife recognizes San Lucas Robin (*T. confinis*) as a separate species, and has assessed each of these two taxonomic concepts independently. To recover some of these IUCN assessments (which, undisputedly relate to imperfectly matched taxon concepts), we downloaded the most recent version of IUCN Red List on 28 November 2024, and incorporated the assessments of 520 species with perfectly matching scientific names in both BirdLife and Clements taxonomies. IUCN assessments are ordinal, ranging from “Least Concern” to “Critically Endangered”. We converted these to a numeric system as outlined in Table 1, and the missing species were imputed following methods detailed below.

**Table 1.** Framework for converting qualitative, ordinal IUCN Red List assessments into quantitative values for downstream analysis. To emphasize the large conceptual leap between species of least concern, which compose 78% of the global avifauna, and species with a more worrisome Red List category, we skip from zero to four when progressing from Least Concern to Vulnerable. Had we not done so, then species considered Vulnerable by the Red List would have been, counterintuitively, ranked closer to those considered Least Concern than those considered Endangered.

| IUCN category | Ordinal score |
| --- | --- |
| Least Concern | 0 |
| Vulnerable | 4 |
| Near Threatened | 5 |
| Endangered | 6 |
| Extinct | 7 |
| Critically Endangered | 7 |
| Extinct in the Wild | 7 |
| Data Deficient | 5 |

The second source of conservation assessment data was the Avian Conservation Assessment Database (ACAD) (Partners in Flight 2024). These are expert-assessed threat levels (Combined Conservation Score, or CCS) that range from 4 to 20 across species, and that are assessed separately for breeding and non-breeding seasons for migratory birds. Of these, we extracted the CCS-max scores from the database, which are, per-species, the higher of the two scores. We used a version of the database we obtained from the Birds of the World on 15 November 2024, which had already been updated to Clements 2024 taxonomy. We converted this to 2023 Clements taxonomy, which resulted in 1,610 species being assigned ACAD scores.

The third and final source of conservation assessment data we used were eBird trend estimates. We obtained these using the *ebirdst* (Strimas-Mackey et al. 2023) package on 24 November 2024. Specifically, we identified all species with available trend estimates, downloaded the data, and derived species-level average, abundance-weighted trends. This was calculated as the sum of the abundance estimate for each 27km^2^ grid cell multiplied by its percent per year trend, divided by the total abundance of the species across all grid cells ((grid cell abundance * ppy)/total abundance across all cells).

### Source of vulnerability factors

We included six separate measures of species’ vulnerability factors, albeit with some redundancy amongst these. Species with small geographic ranges are frequently considered at increased risk of extinction, and it was therefore important to us to include measures of range size. Furthermore, this measure doubles as a proxy for endemism, e.g., to small islands, which is another factor frequently considered in conservation decision making. We included two measures of range size. The first was the area of the BirdLife range maps, i.e., range polygons (Murali et al. 2021). This dataset was available in an older BirdLife taxonomy (10,372 species total in the dataset), and after matching with taxonomic crosswalks available through the AVONET project (Tobias et al. 2022), we were able to retain 9,686 of these range estimates. The second source of range size data we used was based on modeled species ranges from eBird data, again obtained via the *ebirdst* package. Here, we first identified all species with available population status estimates, then downloaded the data at 27km^2^ resolution. For migratory species, we focused only on their breeding season ranges. Then we removed any 27km^2^ cells where the species was not inferred to occur at all, and took the remaining number of cells as an estimate of range size.

Species that are locally rare are also likely at increased risk of extinction, and thus we included a measure of average local abundance as another vulnerability factor. Here, we took the species-level median of the modeled abundance from the eBird breeding and resident grid cells described above (Strimas-Mackey et al. 2023).

It is well documented that some species show a higher tolerance for anthropogenic impacts than others, and we wanted to include a measure of human tolerance in the BirdsPlus species score. A recently published dataset (Marjakangas et al. 2024) provides such an estimate, derived from eBird data and modeled species’ occurrences. We used species’ median “conservative HTI” (human-tolerance index) scores, retaining only those estimates where the scientific name from the dataset directly matched that in Clements 2023 taxonomy (n = 5,985).

Migration has been shown to be the most dangerous part of many species’ life cycles (Rushing et al. 2017; Saunders et al. 2025), and there is mounting evidence that long-distance migrants are faring particularly poorly in today’s world (Rosenberg et al. 2019). As such, it was critical that we include estimates of migration distance. We used two datasets to derive these measures. The first came from (Dufour et al. 2020), specifically the distance_quanti_ALL column, and after taxonomic matching, was available for 10,228 species. Like the first of our range size estimates, these values were derived directly from BirdLife range maps, with added information from reference handbooks (del Hoyo et al. 2017); the column we used corresponds to a migration distance in kilometers for all species, including nominal residents. We further derived our own estimates of migration distance from eBird data. To do so, on 5 May 2023, we downloaded all available eBird data from complete checklists that were neither longer than 12 hours in duration nor 20 km in length. We then grouped these checklists into h3 grid cells (https://h3geo.org/) by month of the year, using a resolution of 6, which corresponds to cells of 36km^2^. Then, per cell-month combination, we found the proportion of total individuals from each of the detected species, and we used these proportions as weights in the following step. Here, per species per month, we identified its geographic centroid as the weighted average of latitude and longitude, after converting to Cartesian coordinates. Finally, we calculated the Haversine distances between each centroid in successive months from January through January of the following year, resulting in 12 distances (usually), which we summed for a final migration distance estimate. We included species which had been detected in at least 10 months of the year, and did not account for any potential movements during these unobserved months (most such data-limited species are tropical and show limited seasonal movements). This resulted in 9,559 species with available eBird-based migration distance estimates.

### Source of phylogenetic and functional uniqueness factors

A species’ independent evolutionary history—the amount of time it has been evolving independently from extant relatives—is another factor that has been suggested as a prioritization tool for conservation (Isaac et al. 2007). We used the phylogeny to calculate species’ edge scores (but without incorporating IUCN status in the calculation) as our measure of “global” phylogenetic uniqueness (n = 11,017). In truth, phylogeny is often incorporated in metrics such as this because it can be a proxy for actual functional diversity and divergence. In our case, however, we were also able to leverage the AVONET dataset (Tobias et al. 2022), a comprehensive morphological database of the worlds’ birds. To derive a measure of global functional uniqueness from morphology, we log-transformed species-level values for: beak length (both from the mechanical hinge as well as from the nares), width, and depth; tarsus length; primary wing length; secondary wing length; tail length; and mass. We then ran a principal components analysis after scaling and centering. Finally, we found the Mahalanobis distance between each species’ point in multivariate space and the centroid of that total space, after accounting for the covariance among the points therein (n = 10,349 species).

We also derived measures of phylogenetic and functional uniqueness that reflect a species’ role in the communities it occurs in. This can be thought of as a coarse measure of phylogenetic or ecological redundancy, where species with large values are unique in comparison to those around them. To derive these measures, we lumped the raw eBird data we downloaded for migration distance calculations into h3 grid cells of 36km^2^. For each species *i*, we identified all grid cells *j* in which it was observed. We excluded grid cells with fewer than three species reported. In each remaining grid cell, we computed the weighted mean distance (phylogenetic or morphological) between species *i* and all co-occurring species *k* ≠ *i*, where the weights were the number of checklists in which each species *k* was reported within that grid cell. Then, for species *i*, we computed a final weighted average across all included grid cells, where the weights were the proportion of checklists in each grid cell *j* in which species *i* was reported. Weights at both steps reflected local species presence (for co-occurring species *k*) and focal species frequency (for species *i* across cells), which ensured that both community context and reporting effort were accounted for. While to our knowledge this is an entirely novel approach, it shares close similarities both to cell-level mean pairwise phylogenetic and functional distance approaches (Miller et al. 2016), and to species-level “fields” (Villalobos et al. 2013). We used these final species-level weighted values as measures of regional phylogenetic and functional uniqueness (n = 10,564 species).

### Normalizing data

To reduce the impact of outliers on the BirdsPlus species scores, and particularly to improve the performance of data imputation procedures described below, we log transformed range size, abundance, migration distance, global phylogenetic and functional uniqueness scores, and regional morphological uniqueness to better approximate normal distributions.

### Phylogenetic imputation

We used *Rphylopars* (Goolsby et al. 2017) to impute missing data. Prior to imputation, every species had at least one known value, and the 25%, 50%, and 75% quantiles for missing columns per species were 4, 4, and 5, respectively (out of 22 columns total). We used an Ornstein-Uhlenbeck model of evolution for the data imputation procedure. This model allows variance to accumulate with random drift, but includes a term to pull the imputed traits back towards the mean. The structure of the phylogeny (specifically the variance-covariance matrix) is used to determine how closely missing species’ traits match their known ancestors’.

### Aggregation into a final BirdsPlus species score

After imputing missing data, we scaled all measures to range from 0 to 1, and then inverted scores for trend, range size, human tolerance, and abundance, such that after inversion species with scores of 1 for these measures were of the greatest conservation concern. Then we established weights for each factor (conservation status, vulnerability factors and uniqueness), summarized in Table 2, and used these to derive a species-level weighted average for each factor. Importantly, however, for measures which were imputed, we multiplied the nominal weight by 0.05, such that it only contributed 5% of its “ideal” weight from Table 2 towards a given factor. Finally, per-species, we summed each of the three factor scores to arrive at a total BirdsPlus species score.

**Table 2.**
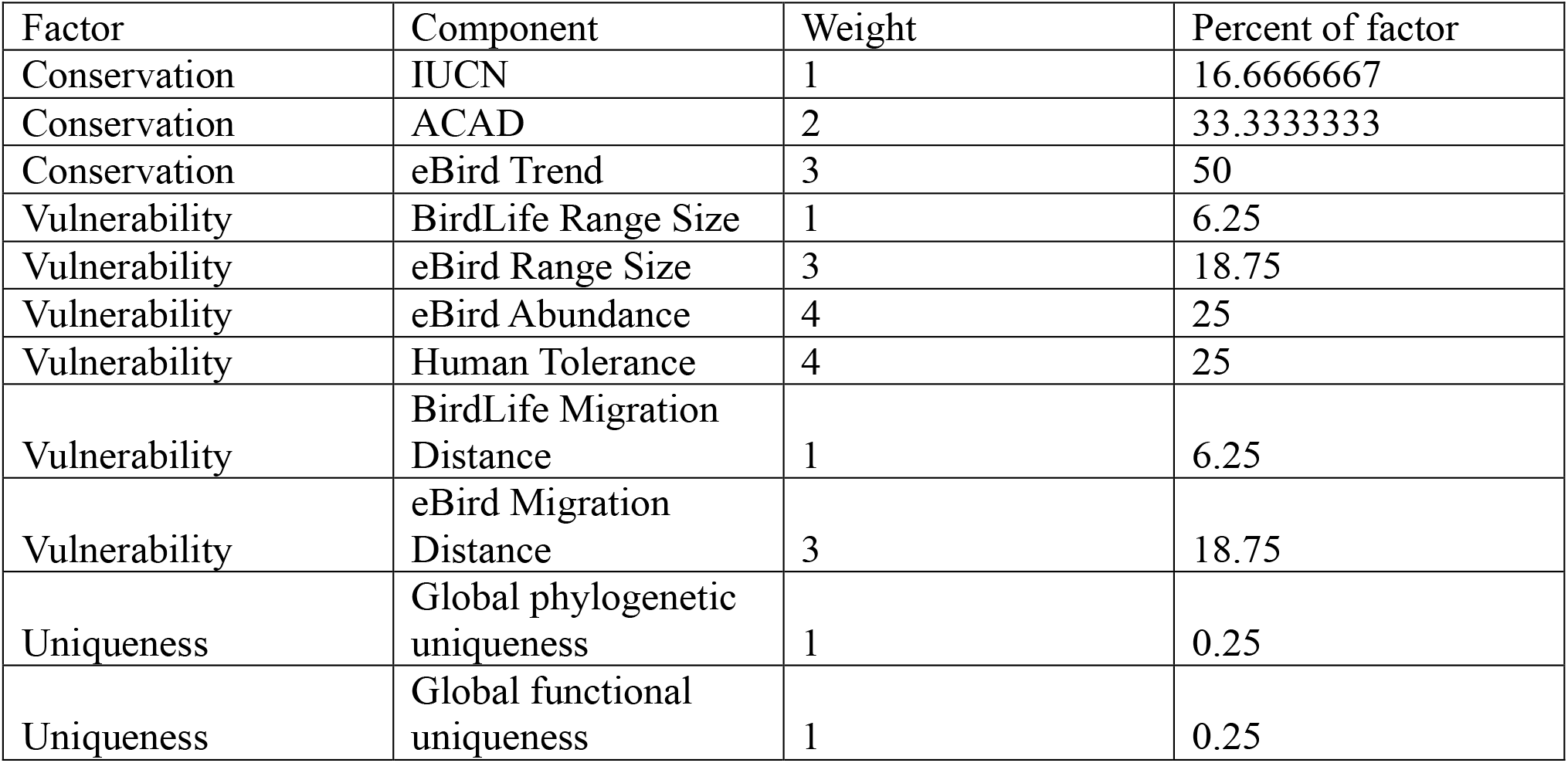
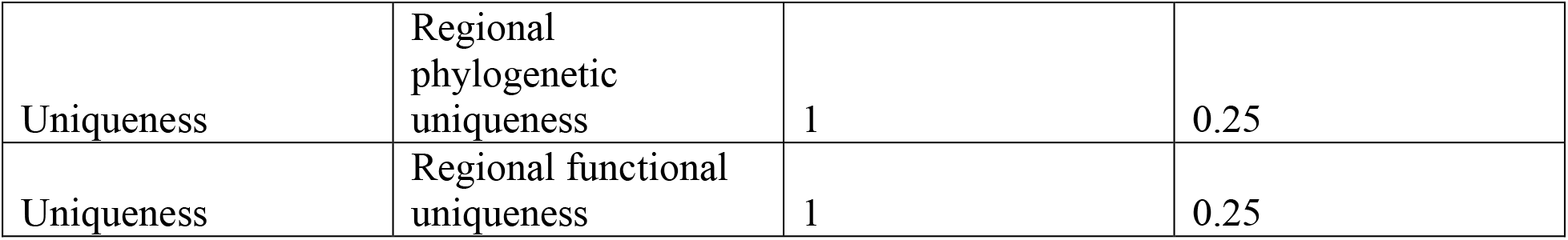
Values used in the weighted averaging step taken to arrive at final conservation, vulnerability, and uniqueness factor scores for each species. These weights are used to defer to preferred databases when the species in question is included in the database. For example, if an eBird trend is available for a given species, 50% of its final conservation score will be driven by its trend as quantified there. However, if the species in question is not included in the database, these weights are multiplied by 0.05, such that the phylogenetically imputed value only imparts 5% of its ideal weight to the final average.

## RESULTS

We derived complete BirdsPlus scores for all 11,017 species of birds in the 2023 Clements taxonomy (Fig. 1, Table S1). These scores are normally distributed and range between 0.58 and 2.72. Depending on the application, users may wish to transform these scores, for example by cubic transformation, to place particular emphasis on species of conservation concern. The code, raw data, and full, imputed dataset are all freely available at the links indicated at the end of the manuscript.

**Figure 1.**
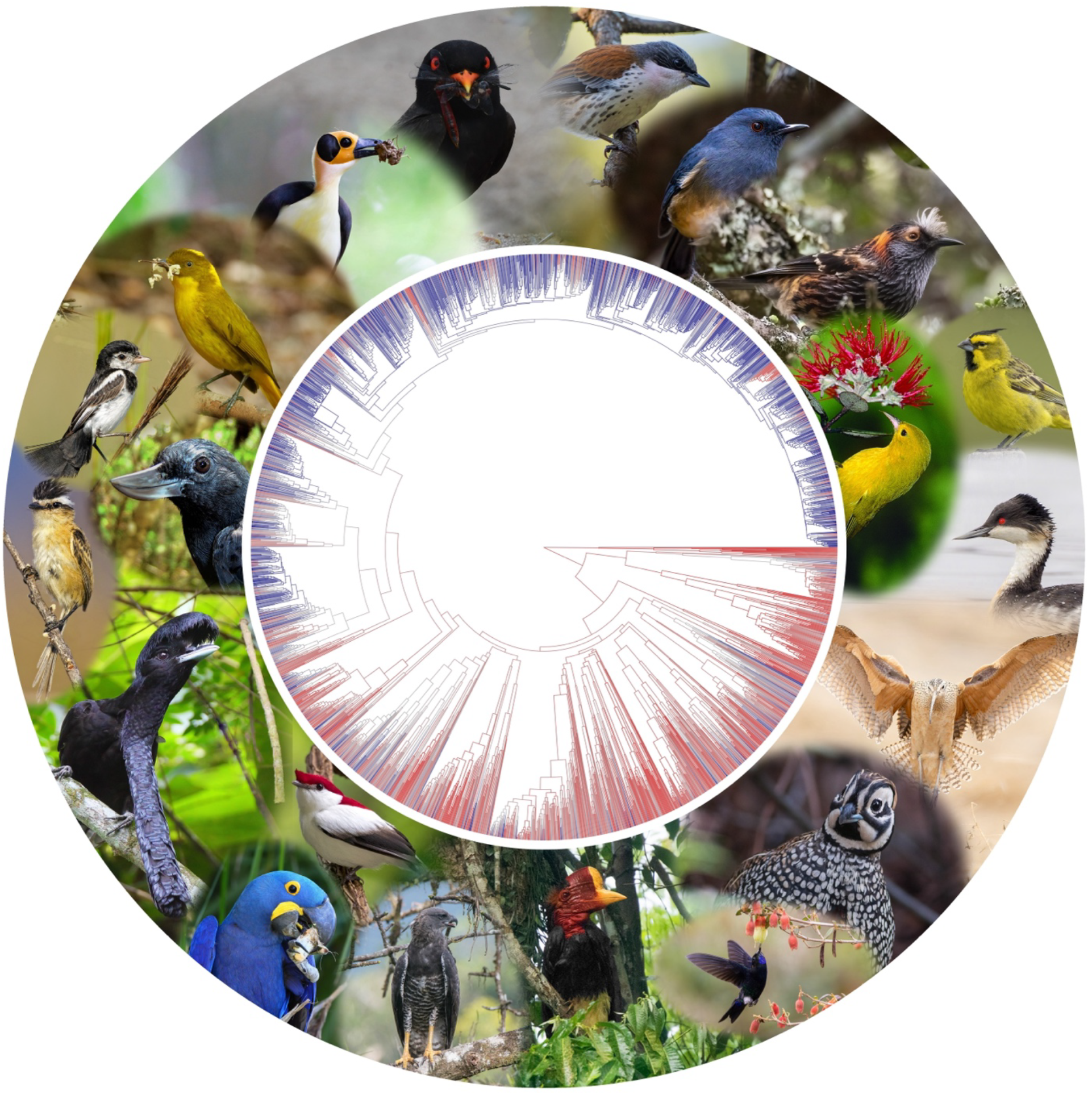
Genus-level average BirdsPlus scores illustrated on a phylogeny of the world’s birds, where genera of greater than average conservation concern, with larger BirdsPlus scores, are colored in red, and those with smaller scores are colored in blue. Clades of well-known conservation concern (including worrisome population statuses, exposure to vulnerability factors, or high ecological and evolutionary uniqueness) are clearly discernible here. Starting at three o’clock, examples include: some grebes, such as Junin Grebe, *Podiceps taczanowskii* (copyright Andrew Spencer, Macaulay Library 382046181); many shorebirds, including Long-billed Curlew, *Numenius americanus* (copyright Jeff Dyck, Macaulay Library 561412711); many Galliformes, including Montezuma Quail, *Cyrtonyx montezumae* (copyright Doug Gochfeld, Macaulay Library 112164941); a number of hummingbirds, including Black-breasted Puffleg and many other *Eriocnemis, E. nigrivestris* (copyright Jay McGowan, Macaulay Library 611304791); most hornbills, including the Helmeted Hornbill, *Rhinoplax vigil* (copyright Ayuwat Jearwattanakanok, Macaulay Library 172593861); some raptors, such as the Crested Eagle, *Morphnus guianensis* (copyright Carlos Bran); a great many parrots (a clade that has also suffered more than its share of extinctions already), particularly macaws like the Hyacinth Macaw, *Anodorhynchus hyacinthinus* (copyright Greg Homel); a fair number of suboscine passerines, including Araripe Manakin, *Antilophia bokermanni* (copyright Ciro Albano), Long-wattled Umbrellabird, *Cephalopterus penduliger* (copyright Greg Homel), Sharp-tailed Tyrant, *Culicivora caudacuta* (copyright Thiago del Toledo e Silva), Recurve-billed Bushbird, *Clytoctantes alixii* (copyright Fundación ProAves), Cock-tailed Tyrant, *Alectrurus tricolor* (copyright Jose Carlos Motta-Junior); some bowerbirds, such as Golden Bowerbird, *Prionodura newtoniana* (copyright Jenny Stiles, Macaulay Library 493305361); a number of odd-looking, species-poor Old World families, including Picathartidae like the White-necked Rockfowl, *Picathartes gymnocephalus* (copyright Daniel López-Velasco | Ornis Birding Expeditions, Macaulay Library 425480471); the two species of enigmatic river martins (one presumed extinct), including African River Martin, *Pseudochelidon eurystomina* (copyright Lionel Sineux, Macaulay Library 627667036); the genus *Laniellus*, which includes Gray-crowned Crocias, *Laniellus langbianis* (copyright Ngoc Sam Thuong Dang, Macaulay Library 399374851); the genus *Sholicola*, including Nilgiri Sholikili, *Sholicola major* (copyright Honza Grünwald, Macaulay Library 620583993); nearly all of the Hawaiian honeycreepers (most already extinct), including Akohekohe, *Palmeria dolei* (copyright Jim Denny) and Anianiau, *Magumma parva* (copyright Peter LaTourrette); and some nine-primaried oscine passerines, especially those targeted for the cage bird trade like Yellow Cardinal, *Gubernatrix cristata* (copyright Dubi Shapiro, Macaulay Library 620559958).

While most of the constituent measures have been published elsewhere, we developed and used three entirely novel measures. Our new, data-driven method of calculating migration distance is available for 9,559 species. The longest distance migrants are all seabirds, while the vast majority of species migrate very little if at all. There is a strong correspondence between our new measure of migration distance and a previously published measure (r^2^ = 0.48, p < 0.001 (Dufour et al. 2020)).

Our regional phylogenetic and functional uniqueness values revealed that species which are closely related to those they co-occur with include members of the families Pellorneidae, Leiothrichidae, and Timaliidae, as well as some seabirds like Pycroft’s Petrel (*Pterodroma pycrofti*). In contrast, those which are phylogenetically unlike others they co-occur with tend to be ratites such as kiwis (*Apteryx* spp.) and tinamous (Tinamidae). Similarly, species which are morphologically akin to others they co-occur with tend to be seabirds, whereas the most functionally irreplaceable species are such things as kiwis, hornbills (Bucerotidae), Shoebill (*Balaeniceps rex*), pelicans (Pelecanidae), large storks (Ciconiidae), and unique island endemics such as Mayr’s Swiftlet (*Aerodramus orientalis*), Inagua Woodstar (*Nesophlox lyrura*) and other Caribbean Trochilinae, and the Vampire Ground-Finch (*Geospiza septentrionalis*). These regional uniqueness values, as well as the h3 grid cell bird assemblages used to derive them, are included as supplemental materials.

## DISCUSSION

Here, we developed an approach for integrating a broad set of factors, from ecological function to current population status, into a unified, continuous score intended to summarize a species’ conservation value in a single number. Conservation practitioners, researchers, policy makers, and funders alike have a need for such assessments. Decades of use of the IUCN Red List for everything from reserve prioritization (Mair et al. 2021) to incorporation in emerging global biodiversity frameworks (Burgess et al. 2024) corroborate this need for additional data-driven methods to guide conservation. Yet, Red List assessments are coarse and ordinal (as opposed to continuous), which limits some downstream uses. Regionally focused and more detailed conservation assessments have been produced, and a variety of other factors of conservation importance can be readily assembled from available databases, but a global project of this scale is currently lacking. Our approach is transparent and dynamic, and while BirdsPlus species scores also reflect our own scientific expertise and judgement, scores are open to easy modification by others; they can be easily recalculated as new information becomes available and taxonomy changes over time.

Our primary purpose in developing these scores has been to provide a quantitative means of valuing site-level biodiversity in a more meaningful way than simple species richness. For example, while the number of species detected from a single point in secondary tropical forest can rival that of primary forests, the species inhabiting these secondary habitats tend to be ecological generalists (Hughes et al. 2020), tolerant of human disturbance (Powell et al. 2015), and to have more secure populations than those inhabiting primary forests. As a result, in order to understand the improved outcomes for biodiversity after conservation management, practitioners often focus on indicator or target species (Kessler et al. 2011). This is certainly a viable approach, but ours is intended to provide an alternative, more comprehensive, and more granular data-driven means of quantifying improved outcomes such as those seen in shade-grown versus sun-grown coffee plantations, for example.

To illustrate how BirdsPlus species scores behave across species with differing ecological roles and a range of conservation statuses, we explore some comparative examples here. Cerulean Warbler (*Setophaga cerulea*), a declining, long-distance Neotropical migrant, has a final BirdsPlus score of 1.47, which is in the 81^st^ percentile of all birds globally. In comparison, the closely related Tropical Parula (*S. pitiayumi*), a largely resident tropical species with a much larger breeding range that co-occurs broadly in the non-breeding season with Cerulean Warbler, has a final score of 0.97, which is in the 10^th^ percentile of all species globally. Cubing these scores, which emphasizes the significance of species of conservation concern (i.e. intentionally introducing some skewness to an otherwise normal distribution), yields scores of 3.15 and 0.91, respectively. Such transformations may be desirable in many cases. At any rate, these differences in final scores are driven in particular by Cerulean Warbler’s increased vulnerability to human impact—the species ranks in the 92^nd^ percentile of all species in terms of migration distance.

Cerulean Warbler also has an elevated conservation status, and it is considered near threatened by the IUCN, has an ACAD score of 14 (as compared to Tropical Parula’s 10), and is declining across its range, whereas Tropical Parula is generally increasing throughout its range, particularly in Mexico (Fink et al. 2023; Strimas-Mackey et al. 2023). As another comparison, while there are many Neotropical species with higher BirdsPlus scores (Tachira Antpitta, *Grallaria chthonia*, for example), when focusing only on those with little to no imputed inputs, the highest ranked is Montezuma Quail (*Cyrtonyx montezumae*, 1.87), while the lowest ranked are Yellow-winged Tanager (*Thraupis abbas*, 0.71) and Palm Tanager (*T. palmarum*, 0.76). Although classified as least concern by the IUCN, Montezuma Quail is a locally uncommon species native to perennial grasslands and oak woodlands of southwestern United States and northern Mexico—plant communities that are routinely overgrazed and also subject to increasing drought—and where it is often hunted and is steeply declining (Stromberg et al. 2020; Fink et al. 2021). In contrast, Palm Tanger (and the closely related Yellow-winged Tanager) is widespread, often found in heavily disturbed environments, and accordingly increasing throughout most of its range (Hilty 2020; Fink et al. 2021); it has a low ACAD score and is ecologically and phylogenetically similar to many other species it co-occurs with.

There is a reasonably close correlation between our new measure of migration distance and a previously published measure (Dufour et al. 2020), as well as tracking data for specific species, e.g., Cerulean Warbler (Raybuck et al. 2022), although neither is a real measure of individual-based migration distance as can be derived from more direct tracking approaches. Notably, our new measure, which does not treat a species’ range as an evenly filled polygon and which uses monthly measurements as opposed to coarse seasonal shifts, appears to address some shortcomings in the range map approach. For example, the three species with the largest positive residuals from a model of the new migration measure as a function of the previous are Peregrine Falcon (*Falco peregrinus*), Blue Seedeater (*Amaurospiza concolor*), and Salmon-crested Cockatoo (*Cacatua moluccensis*), all of which have a migration distance of 0 in the previous measure, but which are inferred to migrate fairly large distances with our new measure. Because some populations of Peregrine Falcon are well known to migrate large distances, and the Blue Seedeater is a nomadic bamboo specialist assumed to move seasonally (García et al. 2023), these shifts suggest increased resolution with our new measure. Salmon-crested Cockatoo is an IUCN vulnerable species restricted in its native range to the South Moluccas, but escaped cage birds are present in Singapore and Hawaii, amongst other regions, and recent changes in eBird now allow these to be reported; this change presumably affected our results, and in the future, we will exclude escapees from the analysis. The three species with the largest negative residuals from the model are Coiba Spinetail (*Cranioleuca dissita*), Arfak Catbird (*Ailuroedus arfakianus*), and Perija Antpitta (*Grallaria saltuensis*), all of which have limited geographic ranges and do not migrate any substantial distances according either to the literature (Billerman et al. 2020) or to our new measure, but which have fairly large migration distances according to the previous measure. Again, these shifts suggest that our new migration values are a significant improvement on previously available measures.

Potential future developments with BirdsPlus scores that warrant additional investigation include incorporation of measures of ecological and cultural significance. Certain species like the Greater Adjutant (*Leptoptilos dubius*, (Barman et al. 2020)) and Philippine Eagle (*Pithecophaga jefferyi*, (Panopio et al. 2021)), for example, carry tremendous cultural significance, and it may be worthwhile to provide a quantitative boost to convey such significance. That said, in these specific cases at least, both species already have BirdsPlus scores in the 96^th^-99^th^ percentile of all species globally. Relatedly, certain species, such as those that excavate nesting cavities (Bednarz et al. 2004) or lead mixed flocks (Zou et al. 2018), offer ecosystem services that may go above and beyond the values currently captured by the evolutionary and functional uniqueness aspects of BirdsPlus species scores, and it might be desirable to capture this significance in the scores in the future. Comprehensive trait databases that convey this significance in a quantitative and continuous way, however, are generally lacking, though some progress has been made (Schuetz & Johnston 2019; Mittermeier et al. 2021). Other planned future developments include the incorporation of additional regional assessments, such as the State of India’s Birds (SoIB 2023) and, ideally, regional variation in species scores, such that invasive species could be downweighted outside their native ranges, and key target species could be upweighted where restoration efforts have been implemented to help conserve them. Last, continuing to shift from broad-stroke IUCN range maps, which frequently overestimate range size (Ramesh et al. 2017), to those that account for actual area of habitat (Brooks et al. 2019), or modeled occurrence when data permits (Fink et al. 2021), is a central goal of our approach going forward.

BirdsPlus species scores (Table S1) provide a simple and intuitive way of conveying the conservation and ecological significance of each of the world’s birds. While there is no doubt conservation is a multidimensional process, and no single score will ever perfectly capture the variety of relevant considerations, there is value in synthesizing existing databases to more accurately reflect the breadth of factors that go into prioritizing species and regions. Simply put, global assessments such as the IUCN Red List have a proven demand, and are rapidly finding their way into globally significant monitoring and verification methodologies (Burgess et al. 2024), but in practice there is extremely limited resolution in these rankings, such as in applications on working lands that might not harbor globally threatened species but provide critical habitat for regionally important species. BirdsPlus scores build upon these assessments by incorporating additional, more detailed regional assessments and factors such as ecological uniqueness and vulnerability factors to arrive at a single score that should be of use both in similar applications to Red List assessments, and particularly to emerging metrics intended to measure progress towards global biodiversity targets.

## Supporting information

BirdsPlus species scores and known and imputed component values

Non-imputed component scores (informed by real data)

Weights used in creating final BirdsPlus species scores

## SUPPLEMENTARY MATERIALS

Table S1. Complete BirdsPlus scores, including all constituent values, for the world’s bird species. These include both real and phylogenetically imputed values. As described in the main text, all values are scaled from 0 to 1 before taking a weighted average using the weights provided in Table 2. Those values which have been imputed can be determined from Table S2.

Table S2. Raw, untransformed values used to derive species’ final BirdsPlus scores. If a cell contains an NA, then the relevant database did not contain a value for that taxon, and the final value in Table S1 was derived via phylogenetic imputation.

Table S3. The final weights used, per species, to derive weighted average conservation status, vulnerability factor, and ecological and evolutionary uniqueness values. Ideal weights are provided in Table 2, but if values were unavailable (see Table S2), these weights have been multiplied by 0.05 to decrease their influence on the final factor-level averages.

